# A deep learning algorithm to translate and classify cardiac electrophysiology: From induced pluripotent stem cell-derived cardiomyocytes to adult cardiac cells

**DOI:** 10.1101/2020.09.28.317461

**Authors:** Parya Aghasafari, Pei-Chi Yang, Divya C. Kernik, Kauho Sakamoto, Yasunari Kanda, Junko Kurokawa, Igor Vorobyov, Colleen E. Clancy

## Abstract

The development of induced pluripotent stem cell-derived cardiomyocytes (iPSC-CMs) has been a critical in vitro advance in the study of patient-specific physiology, pathophysiology and pharmacology. We designed a new deep learning multitask network approach intended to address the low throughput, high variability and immature phenotype of the iPSC-CM platform. It was trained using simulated action potential (AP) data and applied to classify cells into the drug-free and drugged categories and to predict the impact of electrophysiological perturbation across the continuum of aging from the immature iPSC-CMs to the adult ventricular myocytes. The phase of the AP extremely sensitive to perturbation due to a steep rise of the membrane resistance was found to contain the key information required for successful network multitasking. We also demonstrated successful translation of both experimental and simulated iPSC-CM AP data validating our network by prediction of experimental drug-induced effects on adult cardiomyocyte APs by the latter.

## Introduction

The development of novel technologies has resulted in new ways to study cardiac function and rhythm disorders [1]. One such technology is the induced pluripotent stem cell-derived cardiomyocyte (iPSC-CMs) *in vitro* model system [2]. The iPSC-CM system constitutes a powerful *in vitro* tool for preclinical assessment of cardiac electrophysiological impact and drug safety liabilities in a human physiological context [3-8]. Moreover, because iPSC-CMs can be cultured from patient specific-cells, it has shown to be an ideal model system for patient-based medicine [8-10].

While utilization of *in vitro* iPSC-CMs allows for testing of responses to drugs and understanding physiological mechanisms [11-14], there is still a major inherent limitation of the approach: The complex differentiation process to create iPSC-CMs results in a model of cardiac electrical behavior that resembles fetal cardiomyocytes. Hallmarks of the immature phenotype include spontaneous beating, immature calcium handling, presence of developmental currents, and significant differences in the relative contributions of repolarizing potassium currents compared to adult cardiomyocytes (adult-CMs) [15-17]. The profound differences between the immature iPSC-CMs and the adult-CMs have led to persistent questions about the utility and applicability of the iPSC-CM action potential (AP) to predict relevant drug impacts on adult human electrophysiology [18, 19].

Several recent studies have proposed computational frameworks to address the primary limitation in using iPSC-CMs and animal cardiomyocytes for drug screening [11, 12, 20, 21]. The innovative studies described by Tvieto and colleagues [9, 10] presented a translation algorithm that identified a mapping function to identify the relationships between the parameters that are defined by key ion channel conductances in the iPSC-CM APs and the adult-CM APs. In another study by Gong and Sobie, additional insights were revealed through application of an efficient partial least squares regression (PLSR) methodology to translate key physiological features between iPSC-CMs and adult-CMs. They also demonstrated the potential to translate between species, between drug-free and simple drugged models as well as between healthy and diseased phenotypes [20]. Koivumäki et al. also tried to address the problem of iPSC-CMs immaturity by establishing a novel *in silico* mathematical model for iPSC-CMs, which can estimate adult-CM behavior [22].

The efficacy of the linear translation algorithms used in the earlier studies relies on a collection of underlying assumptions [20]. One described by Tvieto et al. is that cardiac protein expression levels would differ but their functional properties remain invariant during maturation and that a drug will modify protein function in the same way for iPSC-CMs and the adult-CMs [11]. Tvieto et al. also acknowledged the difficulty in minimizing the cost function that measures the differences between the initial and target parameters, which therefore required a brute force search algorithm for minimization. One possible explanation for the difficulty in cost function minimization is that linear translation may not capture the nonlinearities comprising the actual underlying physiological differences [20]. Another underlying assumption with linear translation is the required representation of drug effects as a simple pore-block, modeled as a reduction in the maximal conductance of the channel [11, 20]. The earlier studies employed a biased method in that they rely on *a priori* parameter identification and extraction from voltage and calcium traces to allow feature mapping from immature to mature conditions [11, 20]. Earlier translators must also consider drug-free and drugged conditions independently.

In this study, we describe a deep learning multitask network that simultaneously performs translation and classification of signals from simulated cardiac myocytes for both drug-free and drugged conditions and we demonstrate its utility for translating and predicting experimental data as well. The multitask network is an unbiased approach in that the user does not predefine the important parameters of the system. Rather, the network learns from the data to define important parameter regimes and data ranges. The new approach is indifferent to the underlying form of the models and can translate time series data from any source. Moreover, the deep learning approach accepts non-linearity of the system, makes no assumptions about changes in cardiac protein expression and function during maturation and can successfully translate simple pore block and complex conformation state-dependent channel – drug interaction. The network learns from all of these data sources for robust and successful translation, suggesting broad applicability.

Artificial neural networks are increasingly used to advance personalized medicine [23-27]. Long-short-term-memory (LSTM) based networks, which are capable of learning order dependence in sequence prediction problems [28], have been widely used for cardiac monitoring purposes [29-31]. They have been used to extract important biomarkers from raw ECG signals [32-34] and help clinicians to accurately detect common heart failure biomarkers in ECG screenings [32, 35-39]. LSTM networks, which can catch existing temporal information in the electronic health records (EHR), have been highlighted as the best predictive models using real time data [40]. LSTM based classifiers have also empowered early arrhythmia detection by automatically classifying arrhythmias using ECG features [41-45]. In addition, deep learning algorithms have been employed to predict drug-induced arrhythmogenicity associated with blockade of the delayed rectifier K^+^ channel current (*I*_*Kr*_) in the CMs encoded by human ether-à-go-go-related gene (hERG) [46] for sets of small molecules in drug discovery and screening process [46-51].

Here, we implemented a deep learning LSTM based multitask network to classify iPSC-CM AP traces into drug-free and drugged categories and translate them into adult-CM AP waveforms. To collect robust realistic simulated data for training the multitask network, we paced simulated cardiac myocytes with the addition of a physiological noise current at matching cycle lengths for Kernik *in silico* iPSC-CMs [52] and O’Hara-Rudy *in silico* human adult-CMs [53] to generate a population of drug-free simulated cardiac myocyte data. To ensure that our model could perform for both drug-free and drugged iPSC-CM and adult-CM APs simultaneously, we simulated drugged samples via both a simple drug-induced *I*_*Kr*_ block model of hERG channel conduction, *G*_*Kr*_, reduction by *1-50%* and a complex Markov model of conformation-state dependent *I*_*Kr*_ block in the presence of a clinical concentration, *2*.*72 ng/mL*, of a potent hERG blocking drug dofetilide from our recent study [46]. We evaluated the multitask network performance on a test dataset and showed excellent performance to translate and classify signals in the form of time-resolved AP traces. We performed an ablation study to reveal the most important iPSC-CM AP information for network translation into adult-CM APs by removing iPSC-CM AP values during various time frames (feature ablation). We also explored the importance of individual LSTM network building blocks and how decoupling of the translation and classification tasks affected overall network performance. We then showed how proposed multitask network can be applied even to scarce experimental data, which was also used to validate the model.

In this study we show that developments in iPSC-CM experimental technology and cardiac electrophysiological modeling and simulation of iPSC-CMs can be leveraged for the application of artificial neural networks (ANN) as a universal approximator [54] to find the most accurate mapping function which is capable of learning nonlinear relationships to predict disease phenotype and drug response in cardiac myocytes from immaturity to maturation.

## Results

In this study, we set out to build a multitask network that would perform two distinct tasks: The first task is to classify iPSC-CM APs into drug-free and drugged categories. The second goal is to translate iPSC-CM APs into corresponding adult-CM AP waveforms. To collect the data for training the multitask network, we simulated a population of 208 AP waveforms for both Kernik *in silico* human iPSC-CMs [52] (Figure 1E blue) and O’Hara-Rudy *in silico* human adult-CMs [53] (Figure 1F blue). We ensured consistency across a population of simulated myocytes by applying physiological noise at the matching the cycle lengths into the iPSC-CMs and adult-CMs. The cell variability in each population is intended to represent the individual variability that is observed in a drug-free human population [52, 53, 55]. An average AP trace from the population is shown in Figure 1A for iPSC-CMs and Figure 1B for adult-CMs. In Figure 1 panels C and D, the ionic currents underlying the *in silico* iPSC-CM APs and adult-CM APs show marked differences, one reason for the broadly expressed concerns about the applicability of utilizing immature iPSC-CM APs in the study of human disease and pharmacology. The substantial current differences illustrate the necessity of a generalized approach to perform translation from immature myocytes into mature myocytes. To ensure that our multitask network could perform over a range of conditions and model forms, we simulated drugged iPSC-CM and adult-CM APs via both a simple *I*_*Kr*_ drug block model of *G*_*Kr*_ reduction by *1-50%* (250 samples in Figure 1E, F green) and a complex model of conformation-state dependent *I*_*Kr*_ block in the presence of *2*.*72 ng/mL* dofetilide (300 samples in Figure 1E, F purple). We combined the drug-free and drugged models with simple and complex *I*_*kr*_ block model schemes (758 samples) for training the multitask network. The differences in key parameters, upstroke velocity (V_max_), maximum diastolic potential (MDP) and action potential durations (APD) across the three conditions are tabulated and shown in Figure 1G.

**Figure 1.**
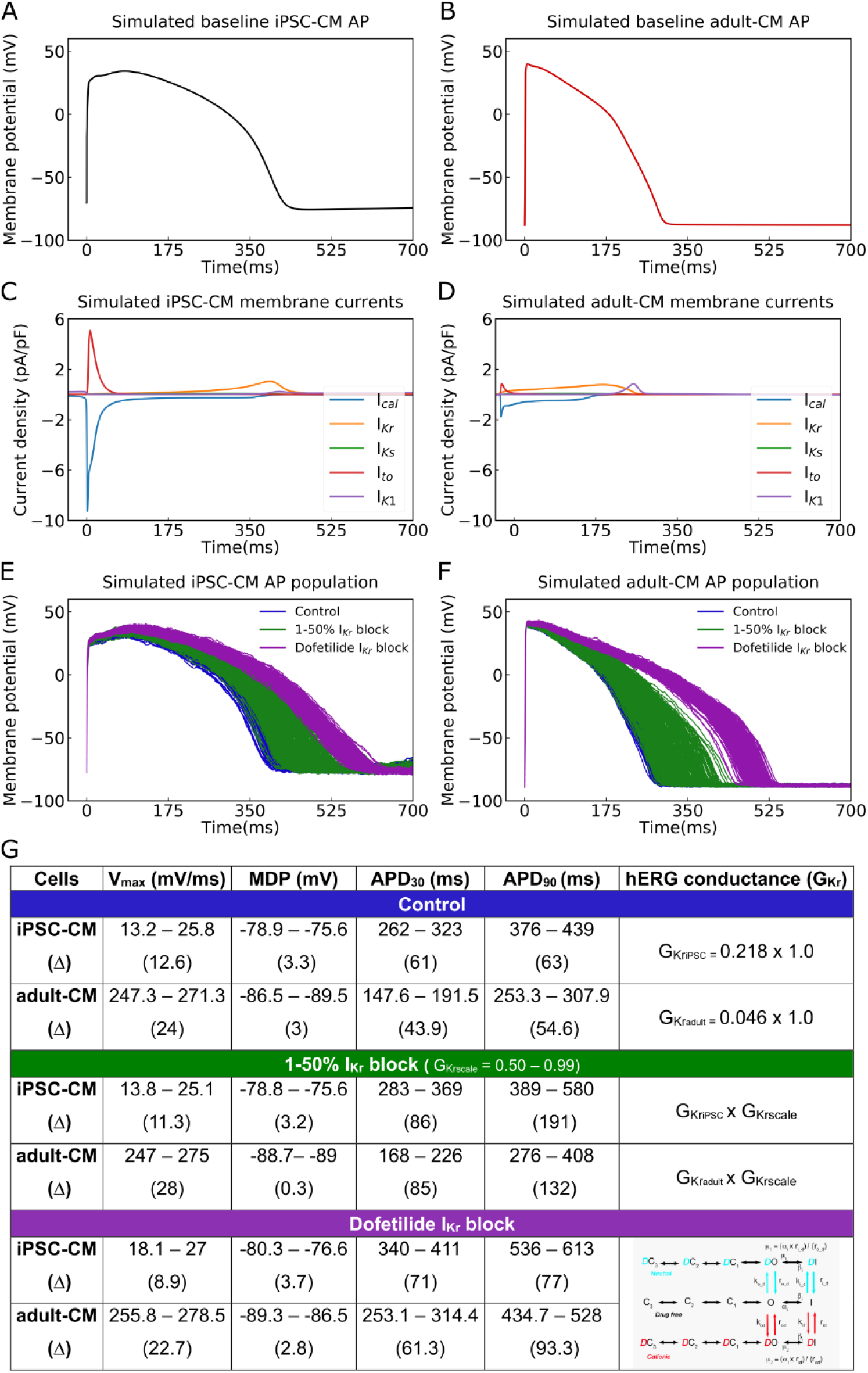
Cellular action potential (AP) and ionic currents for iPSC-CMs and adult-CMs (O’Hara-Rudy human ventricular action potentials). Comparison of Cellular APs in the baseline model of (A) iPSC-CMs and (B) adult-CMs at a matched cycle length of 982 ms. (C – D) Simulated ionic current (*I*_*CaL*_, *I*_*Kr*_, *I*_*Ks*_, *I*_*to*_, *I*_*K1*_) profiles during (C) iPSC-CM and (D) adult-CM APs. (E) APs of spontaneously beating iPSC-CM cells (*n* =208) and (F) adult-CM APs at matched cycle lengths were simulated after incorporating physiological noise currents as drug-free (blue) and drugged *I*_*Kr*_ modeled as simple *G*_*Kr*_ reduction by *1-50% I*_*Kr*_ block (green) and a complex model of conformation-state dependent *I*_Kr_ block in the presence of *2*.*72 ng/mL* dofetilide (purple). (G) Comparison between iPSC-CM and adult-CM drug-free and drugged models with simple and complex *I*_Kr_ block model schemes (as indicated in right column), including upstroke velocity (V_max_), maximum diastolic potential (MDP) and action potential duration (APD).

Next, we applied a digital forward and backward data filtering technique [56] to the simulated iPSC-CM and adult-CM AP traces (Figure 2 left panels). Since we applied physiological noise to introduce a source of variability (as observed in human populations) in our model simulations, we assessed the possible phase distortion for AP waveforms following noise filtering. In Figure 2 (right panels), the distribution of iPSC-CM and adult-CM AP duration at 90% repolarization (APD_90_) values are shown. The near superimposition of the histogram distributions assures that noise filtering does not change the AP waveform morphology or time course and primarily removes existing vertical noises. Panel A and B show simulated drug-free iPSC-CM and adult-CM APs and corresponding APD_90_ distribution with physiological noise in blue and after applying the noise filtering technique in black for iPSC-CM APs and red for adult-CM APs. The same plots are illustrated for drugged AP traces with simple *1-50% I*_*Kr*_ block (Figure 2C and D) and with complex *I*_*Kr*_ block model in the presence of *2*.*72 ng/mL* dofetilide (Figure 2E and F). Next, we normalized drug-free and drugged noise-filtered iPSC-CM APs and adult-CM APs to use them as input and output, respectively, for training the multitask network.

**Figure 2.**
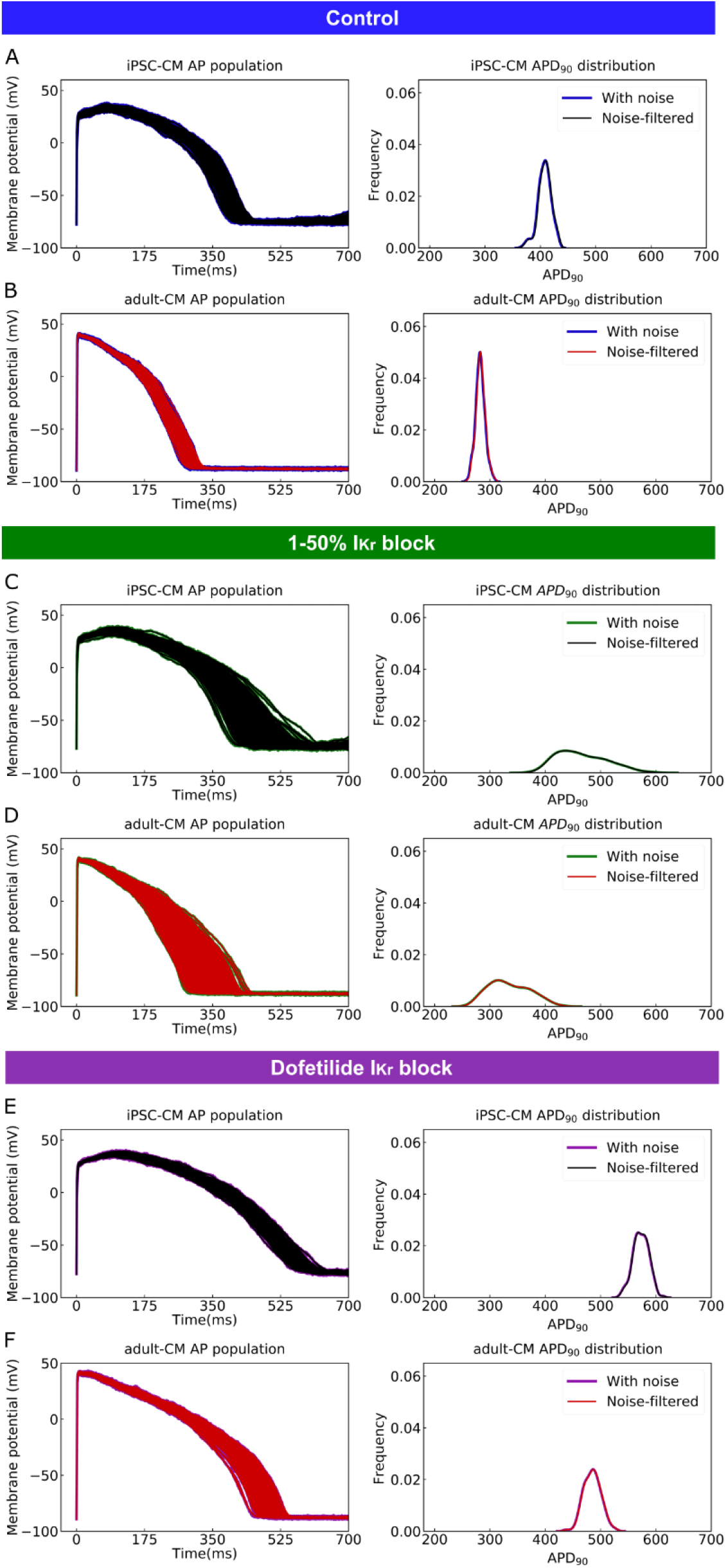
Application of a digital forward and backward data filtering technique to simulated iPSC-CM and adult-CM APs population (left panels) indicates zero phase distortion for APD_90_ value distributions (right panels) for: (A) drug-free iPSC-CM APs with physiological noise in blue and after applying the noise filtering technique in black; (B) drug-free adult-CM APs – blue and red traces; (C) drugged iPSC-CM APs with *1-50% I*_*Kr*_ block – green and black traces; (D) drugged adult-CM APs with *1-50% I*_*Kr*_ block – green and red traces; (E) drugged iPSC-CM APs with *2*.*72 ng/mL* dofetilide – purple and black traces. (F) drugged adult-CM APs with *2*.*72 ng/mL* dofetilide –purple and red traces.

The building blocks of the multitask network are illustrated in Figure 3A. The multitask network receives preprocessed simulation generated iPSC-CM AP waveforms (noise-filtered and normalized) as input and scans whole AP time series values through two stacked LSTM layers (Figure 3A, D). The LSTM layers remember the most important iPSC-CM AP values (features) they need to perform the translation and classification tasks and passes the information to two fully connected layers (Figure 3A, E), one for the translation task to predict the corresponding adult-CM AP waveform (Figure 3B) and one for the classification task to classify iPSC-CM APs into drug-free and drugged categories (Figure 3C).

**Figure 3.**
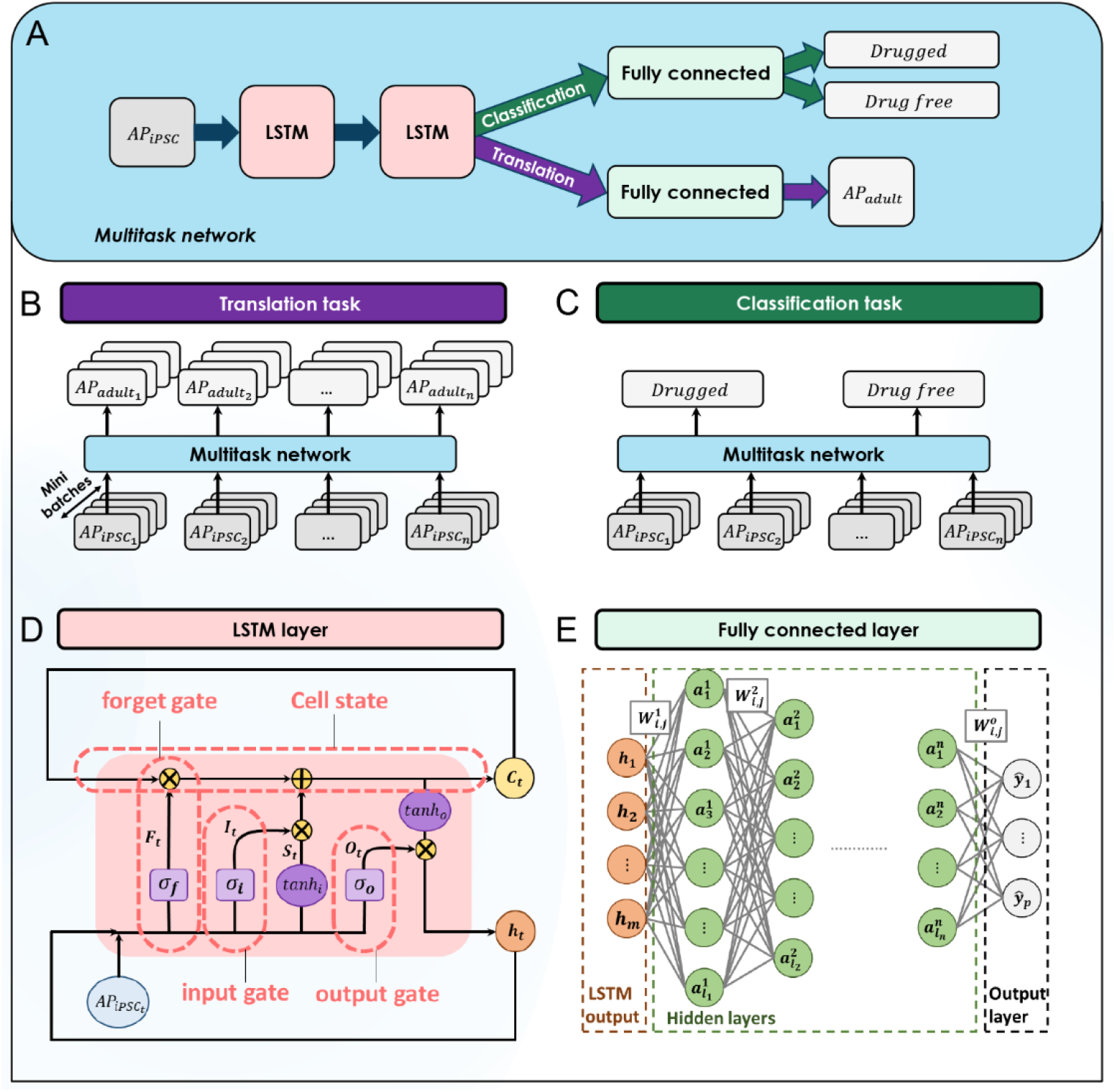
The building blocks of the multitask network. (A) The general overview of the multitask network presented in this study. (B) The translation task to reconstruct adult-CM APs from corresponding iPSC-CM APs. (C) The classification task to classify iPSC-CM APs into drug-free and drugged categories. (D) The logic flow process in the LSTM layers. (E) The architecture of the implemented fully connected layers in the multitask network.

The workflow for training and evaluating the multitask network is depicted in Figure 4. As described above, we generated simulated drug-free and drugged iPSC-CM and adult-CM APs and applied a noise filtering technique to the AP waveforms. The waveforms were then normalized in a data preprocessing step for more efficient training of the multitask network. We used preprocessed iPSC-CM APs as the network input and adult-CM APs along with corresponding drug-free and drugged labels as network outputs, respectively. Next, we randomly split input and output data in 70:10:20 ratio into three subcategories: training, validation, and test data sets. We used the training dataset for training the multitask network to simultaneously perform translation and classification. The mean squared error, R^2^-score [57] and error in adult-CM APD_90_ prediction were used as evaluation metrics for the translation task. For the classification task, area under the receiver operating characteristic (AUROC) curve [58], network prediction accuracy, precision and recall [59] were used to evaluate the network performance. To prevent overfitting, we calculated the evaluation metrics for both tasks using validation data during each iteration of training and compared those with values from the training dataset. When the model performance on the training dataset exhibited degradation relative to the validation dataset, we ceased training and tuning of the network hyperparameters. We evaluated the underlying mechanisms that inform the network performance by using a holdout test data set to perform an ablation study. The ablation study allowed us to identify the most important information for network performance and is an indicator of the data that the network deems most important to remember to allow accurate translation into adult-CM APs (feature ablation). Finally, we performed a type of network component dissection by sequentially eliminating individual LSTM layers or the classification task to determine if all elements of the network are important to the overall performance.

**Figure 4.**
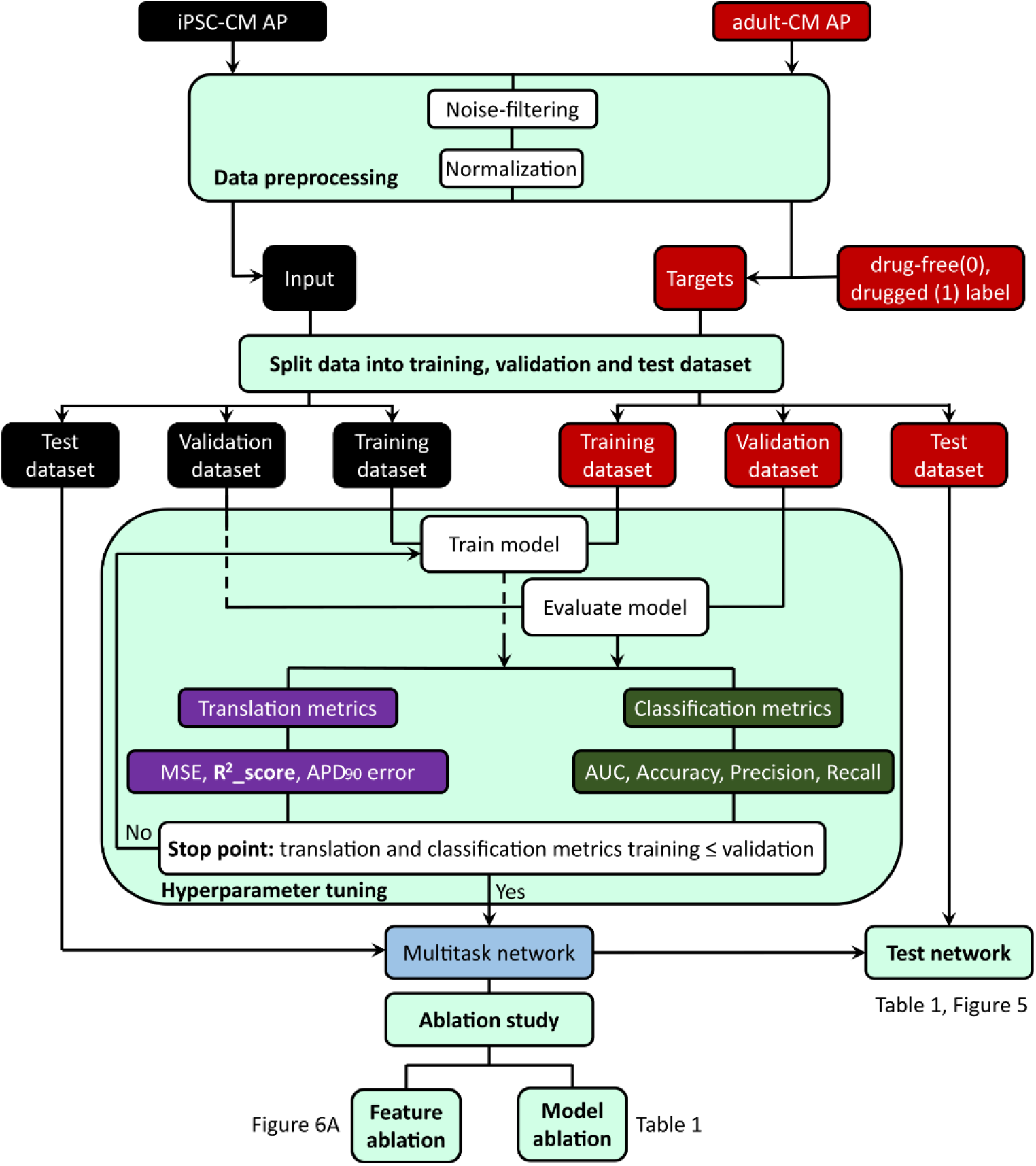
Machine learning workflow in this study: 1) Data preprocessing includes noise-filtering and normalization of the drug-free and drugged iPSC-CM and adult-CM APs; 2) Incorporating the preprocessed iPSC-CM APs as input and adult-CM APs and corresponding labels (drug-free (0) and drugged (1)) of iPSC-CM APs as targets into the multitask network; 3) Splitting the input and target data into training, validation and test set, and using training and validation set for training and tuning the network hyperparameters; 4) Comparing the network performance for training set and validation set to decide when to stop training and tuning the network hyperparameters; 5) Testing the overall multitask network performance using holdout test dataset and removing the LSTM layers, classification task (model ablation) and iPSC-CM AP values at different time frames (feature ablation) to study the performance of the network in the absence of its building blocks.

Figure 5 and Table 1 illustrate the overall multitask network performance for translation and classification tasks for the training and test data sets. Panels A and D in Figure 5 represent iPSC-CM APs (black), which were used for training and testing the multitask network, respectively. Panels B and E depict the comparison between simulated (red) and translated (cyan) adult-CM APs used for the training and testing the network. The comparison between histogram distribution of APD_90_ values for simulated and translated adult-CM APs in Figure 5C and F show good agreement in terms of the frequency of virtual cells with similar APD.

**Table 1.** Statistical measures for evaluating the performance of the multitask network for both iPSC-CM AP trace classification into drug-free and drugged categories and their translation into adult-CM APs for training and test datasets as well as the effect of removing LSTM layers and classification task on the network performance.

| Translation |  |  |  |  |
| --- | --- | --- | --- | --- |
| Performance metrics | MSE | R<sup>2</sup>_score | Error in APD<sub>90</sub> prediction |  |
| *Training dataset* | 0.0027 | 0.992 | 3.41% |  |
| *Test dataset* | 0.0029 | 0.991 | 3.60% |  |
| *Remove LSTM layers test dataset* | 0.0031 | 0.991 | 3.78% |  |
| *Remove classification task test dataset* | 0.0034 | 0.990 | 4.33% |  |
| Classification |  |  |  |  |
| Performance metrics | AUROC | Accuracy | Recall | Precision |
| *Training dataset* | 0.93 | 92% | 0.92 | 0.93 |
| *Test dataset* | 0.91 | 92% | 0.92 | 0.92 |
| *Remove LSTM layers test dataset* | 0.90 | 92% | 0.90 | 0.91 |

**Figure 5.**
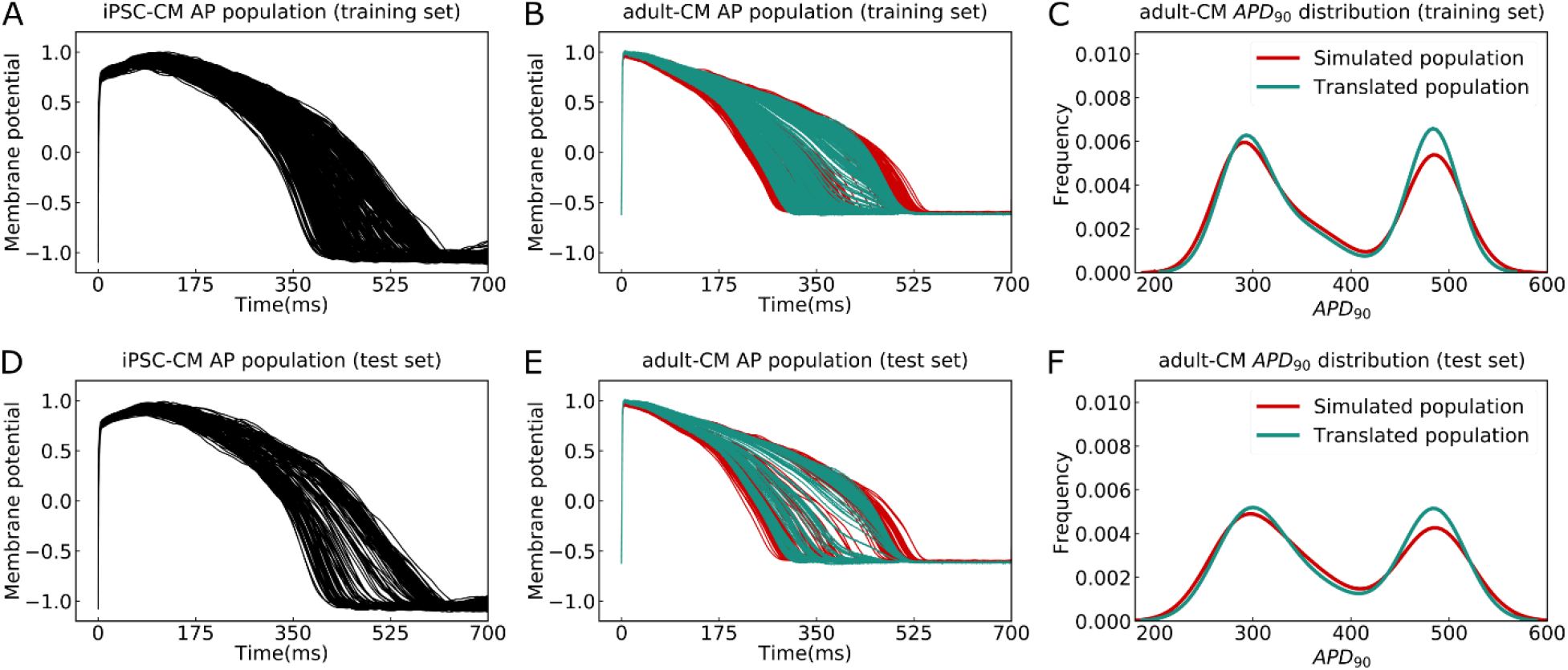
The performance of the multitask network for translating iPSC-CM APs into adult-CM APs. (A) The iPSC-CM APs used for training the multitask network contained a variety of drug-free and drugged action potential morphologies (Training set). (B) Comparison between simulated (red) and translated adult-CM APs (cyan) in the training set. (C) Comparison between the histogram distribution of APD_90_ values for simulated and translated adult-CM APs in the training set. (D) Dedicated iPSC-CM APs for testing the performance of the multitask network (Test set) (E) Comparison between simulated (red) and translated adult-CM APs (cyan) in the test set. (F) Comparison between histogram distribution of APD_90_ values for simulated and translated adult-CM APs in the test set.

The performance evaluation metrics for both the translation and classification tasks are listed in Table 1. The multitask network exhibits high accuracy in performing translation, despite large variability in APDs and regardless of the underlying model form. The network is able to translate iPSC-CM APs into adult-CM APs with less than *0*.*003* mean-squared error (MSE), *0*.*99* R^2^_score and less than *4%* error in APD_90_ prediction for both training and test datasets. To evaluate the network performance for the classification task we compared the AUROC, prediction accuracy, recall and precision for both training and test datasets. The multitask network proved to perform well in categorizing iPSC-CM APs into drug-free and drugged waveforms with approximately *90%* accuracy (Table 1). Finally, we performed a type of network component dissection by sequentially eliminating individual LSTM layers or the classification task to determine if all elements of the network are important to the overall performance. The impact of removing these elements of the network on the network performance is shown in Table 1.

Next, we performed a “computational” ablation study as a correlate to the types of physiological ablations that are used to examine the roles and functions of a physiological system [60, 61]. We tested how the performance of the multitask network would change by removing various information contained within specified time frames as shown in Figure 6A. To reveal the most important iPSC-CM AP information for translation into adult-CM APs, we did not allow the network to process data from within designated time frames from the iPSC-CM APs (feature ablation). We then retrained the multitask network by setting the missing information equal to zero and compared the calculated MSE in adult-CM APs translation (red bars) with the recorded MSE for multitask network (green line) when it was provided full access to the complete iPSC-CM AP data. We observed that network is extremely sensitive to information contained within the *400-500 ms* timeframe (blacked dashed bar in Figure 6A).

**Figure 6.**
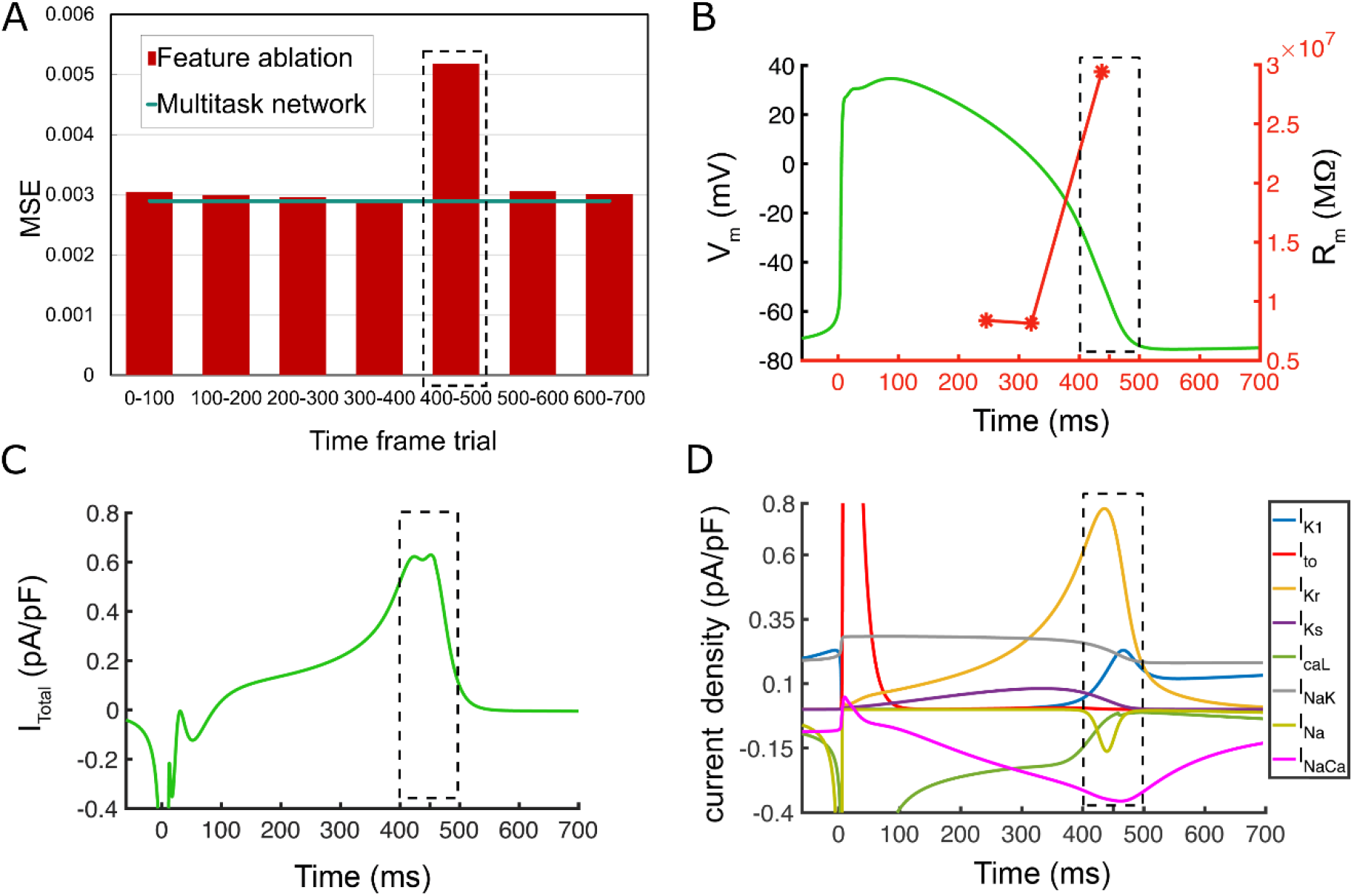
The feature ablation study on the proposed multitask network is performed by removing iPSC-CM AP values during different time frames and evaluating their importance on adult-CM AP translation. The largest effect (most important information) is observed at *400-500 ms* interval (dotted black line). (A) Comparison between intact multitask network MSE (cyan line) and obtained MSE values for adult-CM AP translation during removal of indicated time frames within iPSC-CM APs (red bars). (B) AP trace (green) and membrane resistance (red) as a function of simulation time indicating very high values (as dI **→** 0, dV/dI **→** ∞) for the latter at *400-500 ms*. (C) Total current density, I_total_, demonstrates a plateau followed by a rapid decline at *400-500 ms*. (D) Individual current densities indicate a period of inward and outward current balance followed by rapid changes in *I*_*Kr*_ and other repolarizing components at *400-500 ms* time interval.

This result suggests that the most important information needed to distinguish adult-CM AP signals from iPSC-CM AP signals is contained in a particular region of the AP plateau. The timeframe of the AP between 400 and 500 ms (Figure 6A), corresponds to a phase of exquisite sensitivity to perturbation. We have identified this particular AP range in an earlier study as the phase when the membrane resistance of the myocyte increases markedly (Figure 6B) [62]. This occurs as the inward and outward currents balance each other, leading to a net whole cell current that is nearly constant so that dI **→** 0, dV/dI **→** ∞ (Figure 6C), followed by a rapid reduction in outward current. Figure 6D demonstrates that individual current densities have a period of inward and outward current balance followed by rapid changes in *I*_*Kr*_ and other repolarizing currents at *400-500 ms* time interval.

We next set out to demonstrate the real-world utility of the multitask classification and translation network by applying the network to experimental data. We used experimental iPSC-CM APs from the Kurokawa lab (Figure 7A) as the input data into the multitask network and translated to predicted adult-CM APs as shown in Figure 7B. The translation notably resulted in a reduction in variability in APD in the adult translated cells, consistent with our simulated results and with previous experimental observations [18, 63]. In an additional validation of the multitask network, we undertook a test of the network to accurately translate drug block in iPSC-CMs to adult AP effects and then compared the predicted results with measured experimental data [53]. We first simulated iPSC-CM APs with *50%* block of *I*_*Kr*_. We then used these simulated APs as an input for the multitask network and used the output from the translation task to predict *50%* block on adult-CMs. In Figure 7C, the translated drugged APD_90_ values are shown as turquoise asterisks plotted against simulations from O’Hara-Rudy adult-CM APs with *50% I*_*Kr*_ block (red curve) and experimental *50%* block of *I*_*Kr*_ by *1μM* E-4031 (blue squares) [53]. These data validate that the effects of drug block in iPSC-CMs can be successfully translated to predict its effect on adult human cardiomyocyte APs.

**Figure 7.**
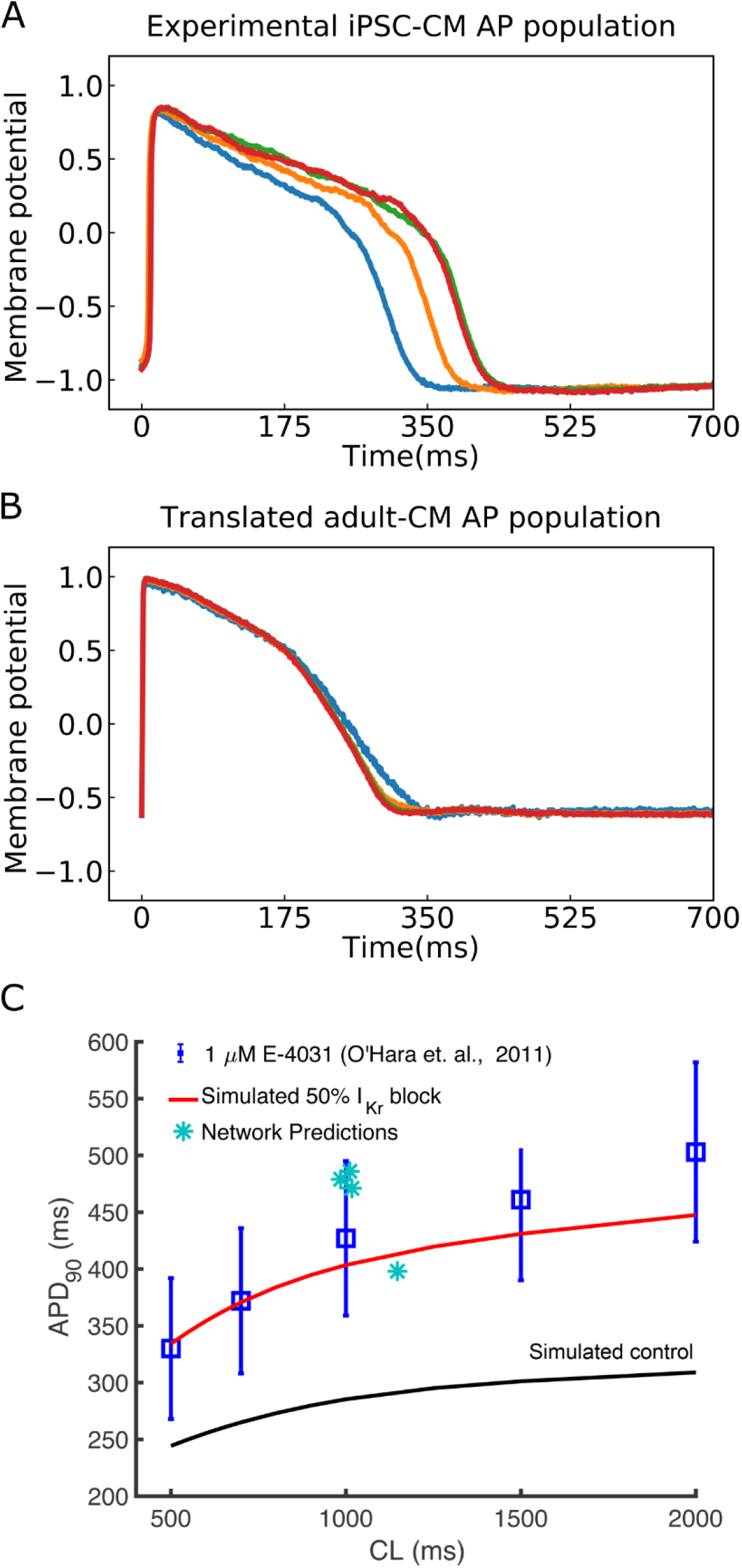
Translation of experimentally recorded iPSC-CM APs into adult-CM APs to validate the multitask network performance. (A) Experimentally recorded iPSC-CM APs from the Kurokawa lab. (B) Translated adult-CM APs from experimentally recorded iPSC-CM APs via the multitask network. (C) Comparing translated adult-CM APD_90_ values with *50% I*_*Kr*_ block (turquoise asterisks) with previously published simulated (red curve for drugged and black for drug-free control) and experimental (blue squares) values from O’Hara-Rudy study [1] indicates model validation.

## Discussion

In this study, we developed a data-driven deep learning approach to address well known shortcomings in the induced pluripotent stem cell-derived cardiomyocyte (iPSC-CM) platform. A concern with iPSC-CM is that the data collection results in measurements from immature action potentials, and it is unclear if these data reliably indicate impact in the adult cardiac environment [14, 64-68]. Here, we set out to demonstrate a new way to allow translation of results from the iPSC-CM to a mature adult cardiac response. The deep learning network also revealed new mechanisms that are critical to convert iPSC-CM APs to mature adult cardiac APs.

Application of a deep learning artificial neural network to simultaneously translate and classify signals from simulated iPSC-CMs for both drug-free and drugged conditions has several key advantages. Because there is no need for the multitask network user to *a priori* define the important system parameters, the approach is by definition an unbiased model. A key part of the “artificial intelligence” is learning from the data to make decisions about which elements of the data are the most important. Another benefit is the model-agnostic approach in that the learning network is indifferent to the underlying form of the models and can readily translate time series data from any source. The non-linearity of the system is accepted by the deep learning approach, and there are no assumptions made about cardiac protein expression levels and changes in their function during cardiomyocyte maturation. The deep learning artificial neural network can successfully translate simple pore block and complex conformation state-dependent channel – drug interaction models. The network can learn from multiple sources of data even when they are generated from different models and learns from all the data sources concurrently for robust and successful translation. All of these aspects of the technology presented here suggest broad applicability for use across ages, species and conditions and we demonstrate its utility for translating and predicting experimental data.

The multitask network presented here performed well in the setting of the noted variability in measurements from iPSC-CM APs. As described in Figure 1, we utilized a modeling and simulation approach from our recent study [52, 69] to generate a population of iPSC-CM action potentials that incorporate variability comparable to that in experimental measurements. Utilizing simulated data presented a unique opportunity: We were able to generate large amounts of data that were used both to train and optimize the network and then to test the network with specifically designated distinct simulated data sets. Utilizing simulated data to train a deep learning network may constitute a widely applicable approach that could be used to train variety of networks to perform multiple functions where access to comparable experimental data is not feasible.

The multitask network exhibits high accuracy in performing translation, despite large variability in APDs and regardless of the underlying model form (Figure 5 and Table 1). The network was able to translate iPSC-CM APs into adult-CM APs with less than *0*.*003* mean-squared error (MSE), *0*.*99* R^2^_score and less than *4%* error in *APD*_*90*_ prediction for both the training and test dataset. To evaluate the network performance for the classification task we compared the AUROC, prediction accuracy, recall and precision for both training and test datasets. The multitask network proved to perform well in categorizing iPSC-CM APs into drug-free and drugged waveforms with approximately *90%* accuracy (Table 1). Finally, we performed a type of network component dissection by sequentially eliminated individual LSTM layers or the classification task to determine if all elements of the network are important to the overall performance. The impact of removing these elements of the network on its performance is shown in Table 1. The studies show that the multi-task network conferred additional benefit over considering the translation task alone. For example, we noted that adding the classification task to distinguish drug-free and drugged action potentials could improve the performance of the translation task (Table 1).

When we performed an ablation study to prevent the deep learning network from using information within prespecified time windows, the results revealed that the most important information needed to predict adult-CM APs from iPSC-CM AP signals is contained in the phase of the AP between 400 and 500 ms (Figure 6). This result suggests that the most important information needed to distinguish iPSC-CM AP signals from adult-CM AP signals is contained in the range of the AP that corresponds to a phase of exquisite sensitivity to perturbation. We have identified this particular AP range in an earlier study as the phase when the membrane resistance of the myocyte increases markedly (Figure 6B) [62].This occurs as the inward and outward currents balance each other, leading to a net whole cell current that is unchanging (dI **→** 0, dV/dI **→** ∞), followed by a rapid reduction in the outward current (Figure 6C and D).

Following the optimization and demonstration of the network as an accurate tool for both translating and classifying data, we then used the same network to translate experimentally obtained data. We showed that the proposed network can effectively take experimental data as an input from immature iPSC-CM APs and translate those data to produce adult action potential waveforms. It is notable that the variation observed in the adult-CM AP duration is smaller compared to iPSC-CM APDs (Figure 7A-B). This has been observed both experimentally [18, 63] and in our simulated cell environment [52, 69]. Although the simulated iPSC-CM has a large initial calcium current (Figure 1C) compared to the simulated adult-CM (Figure 1D), the amplitude of currents flowing through adult-CM action potential plateau is notably larger. The immature iPSC-CM cells have low conductance during the AP plateau rendering it comparably higher resistance. For this reason, small perturbations to the iPSC-CM APs have a larger impact on the resulting AP duration than observed in adult cells [62]. We also used simulated iPSC-CM APs subject to *50%* block of *I*_*Kr*_. We translated those data to adult-CM APs and then compared with the previously reported impact of *50% I*_*Kr*_ block on adult human cell APs from experiments [53] and noted excellent agreement thereby providing validation of our network.

In this study, we show that a deep learning network can be applied to classify cells into the drug-free and drugged categories and can be used to predict the impact of electrophysiological perturbation across the continuum of aging from the immature iPSC-CM action potential to the adult ventricular myocyte action potential. We translated experimental immature APs into mature APs using the proposed network and validated the output of some key model simulations with experimental data. The multitask network in this study was used for translation of iPSC-CMs to adult APs but could be readily extended and applied to translate data across species and classify data from a variety of systems. Also, another extension of the technology presented here is to predict the impact of naturally occurring mutations and other genetic variations [70].

## Methods

### Simulated data for training and testing the multitask network

#### The drug-free iPSC-CM and adult-CM action potentials

The Kernik *in silico* iPSC-CM baseline cells were paced from resting steady-state. The O’Hara-Rudy *in silico* endocardial cell model was used for the baseline adult-CMs [53]. The control adult-CMs were paced at the cycle length of *982 ms* to match the cycle length of the last beat of the spontaneously depolarizing iPSC-CM AP. The iPSC-CM AP populations (*n* =208) were generated by incorporating physiological noise (see **Simulated physiological noise currents** section below). The adult-CMs were paced with noise for *100* beats after reaching steady state at the matching cycle length of the last beat of iPSC-CM AP populations. The numerical method used for updating the voltage was Forward Euler method [71].

#### A simple drug-induced 1-50% I_Kr_ block model through G_Kr_ reduction

The iPSC-CMs and the adult-CMs populations were paced with 1-50% I_Kr_ block with 1% increments. This was accomplished by scaling down hERG channel (*I*_*Kr*_) conduction, *G*_*Kr*_, by the fraction of the block, *G*_*Krscale*_, in the 0.50 – 0.99 range with 0.01 decrements (see central rows in Fig. 1G). The adult-CM model was simulated at five varying beating rates for each percentage of block that matches to the last beat of iPSC-CMs with *1-50% I*_*Kr*_ block (*n* = 250). For example, one drugged adult-CM (*50% I*_*Kr*_ inhibition) was paced at cycle length of *1047 ms* to match the cycle length of the last beat of iPSC-CMs AP with *50% I*_*Kr*_ block.

#### Complex model of conformation-state dependent I_Kr_ block in the presence of 2.72 ng/mL dofetilide

The *I*_*Kr*_ channel Hodgkin-Huxley model in both iPSC-CM and adult-CM AP models was replaced with a drug – hERG channel interaction Markov model (see bottom rows in Fig. 1G) that we have previously published [72]. iPSC-CM (*n* = 300) and adult-CM AP populations (*n* = 300) were generated with physiological noise in the presence of *2*.*72 ng/mL* dofetilide, a potent hERG channel blocker. The adult-CM populations were paced with dofetilide for 100 beats after reaching steady state at the matching cycle length of the last beat of iPSC-CM AP populations with dofetilide as described above.

### Simulated physiological noise currents

Simulated noise current was added to the last 100 paced beats in the simulated AP models, and simulated APs were recorded at the 2000th paced beat in single cells. This noise current was modeled using the equation from [55],

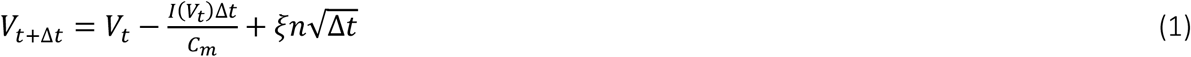

Where *n ϵN*(*0,1*) is a random number from a Gaussian distribution, and *Δt* is the time step. *ξ*; = *0*.*3* is the diffusion coefficient, which is the amplitude of noise. The noise current was generated and applied to membrane potential, *V*_*t*_, throughout the last 100 beats of simulated time course.

### Experimental iPSC-CMs

Human iPSC-CMs (201B7, RIKEN BRC, Tsukuba, Japan) were cultured and subcultured on *SNL76/7* feeder cells as described in detail previously [73]. Cardiomyocyte differentiation was performed as described [73]. Commercially available iCell-cardiomyocytes (FUJIFILM Cellular Dynamics, Inc., Tokyo, Japan) were cultured according to the manual provided from the company. Action potentials were recorded with the perforated configuration of the patch-clamp technique as described in detail previously [73]. Measurements were performed at *36 ± 1 °C* with the external solution composed of (in *mM*): *NaCl* (*135*), *NaH*_2_*PO*_*4*_ (*0*.*33*), *KCl* (*5*.*4*), *CaCl*_*2*_ (*1*.*8*), *MgCl*_*2*_ (*0*.*53*), glucose (*5*.*5*), *HEPES, pH 7*.*4*. To achieve patch perforation (*10-20 MΩ*; series resistances), amphotericin B (*0*.*3-0*.*6 µg/mL*) was added to the internal solution composed of (in *mM*): aspartic acid (*110*), *KCl* (30), *CaCl*_*2*_ (*1*), adenosine-*5’*-triphosphate magnesium salt (*5*), creatine phosphate disodium salt (*5*), *HEPES* (*5*), *EGTA* (*11*), *pH 7*.*25*. In quiescent cardiomyocytes, action potentials were elicited by passing depolarizing current pulses (*2 ms* in duration) of suprathreshold intensity (*120 %* of the minimum input to elicit action potentials) with a frequency at *1 Hz* unless noted otherwise.

### The multitask network architecture

The multitask network was comprised of two stacked LSTM layers followed by independent fully connected layers (Figure 3A) for the classification and translation tasks. The LSTM layers memorized the important information the network needed to perform two discussed tasks and then transferred the extracted information (features) into the subsequent fully connected layers to translate iPSC-CM APs into adult-CM AP waveforms (Figure 3B) and classify iPSC-CM APs into drug-free and drugged categories (Figure 3C).

#### Long-short term memory (LSTM) layers (Figure 3D)

We used LSTM layers as the first two layers of the multitask network to promote network temporal information learning which data in a sequence was important to keep or to throw away. At each time step, the LSTM cell took in three different pieces of information, the current input data 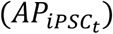, incoming short-term memory (hidden state) (*h*_*t*−1_) and incoming long-term memory (cell state) (*C*_*t*−1_). The LSTM layers were responsible for extracting the most important information while scanning the AP traces using the short- and long-term memory components. The short-term memory weighted the importance of AP values at subsequent time steps and long-term memory has been using the short-term memory to decide the overall importance of all AP values from the beginning (*t* = 0 ms) to the end (*t* = 701 ms) for performing classification and translation tasks. The LSTM cells contained internal mechanisms called gates. The gates were neural network with weights (*w*) and bias terms (*b*) that regulated the flow of information at each time step before passing on the long-term and short-term information to the next cell [74]. These gates are called input gate, forget gate, and output gate (Figure 3D).

The forget gate, as the name implies, determined which information from the long-term memory should be kept or discarded. This was done by multiplying the incoming long-term memory by a forget vector generated by the current input 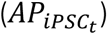 and incoming short-term memory (*h*_*t*−1_). To obtain the forget vector, the incoming short-term memory and current input were passed through a sigmoid function (*σ*_*f*_) [75]. The output vector of sigmoid function, *F*_*t*_, (Eq. 2) was a binary comprising 0s and 1s and was then multiplied by the incoming long-term memory (*C*_*t*−1_). to choose, which parts of the long-term memory were retained.

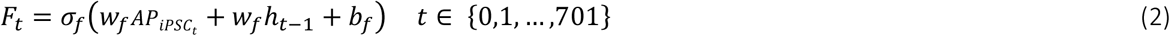

The input gate decided what new information is being stored in current long-term memory (*C*_*t*_). It considered the current input 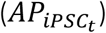 and the incoming short-term memory (*h*_*t*−1_) and transformed the values to be between 0 (unimportant) and 1 (important) using a sigmoid activation function (*σ*_*i*_) (Eq. 3). The second layer in input gate took the incoming short-term memory (*h*_*t*−1_) and current input 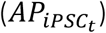 and passed them through a hyperbolic tangent activation function (*tanh*_*i*_) to regulate the network computation (Eq. 4).

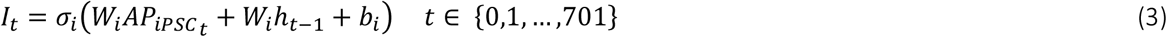

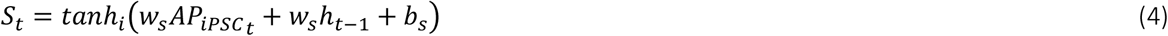

The outputs from the forget and input gates then underwent a pointwise addition to find the current long-term memory (*C*_*t*_) (Eq. 5), which was then passed on to the next cell.

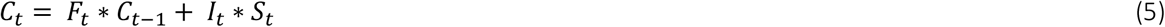

Finally, the output gate utilized current input 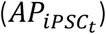 and the incoming short-term memory (*h*_*t*−1_) and passed them into a sigmoid function (*σ*_*0*_) (Eq. 6). Then the current long-term memory (*C*_*t*_) passed through a *tanh* activation function (*tanh*_*0*_) and the outputs from these two processes were multiplied to produce the current short-term memory *h*_*t*_ (Eq. 7).

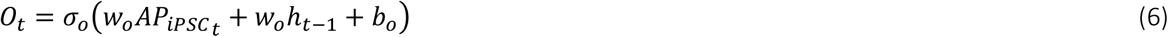

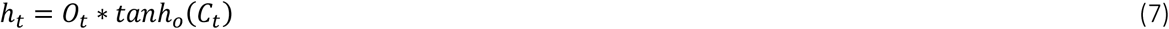

The short-term and long-term memory produced by these gates were carried over to the next cell for the process to be repeated. The output of LSTM layers for each time step (*h*_*t*_) was obtained from the short-term memory, also known as the hidden state, and was subsequently passed into fully connected layers to perform the translation and classification tasks as described below.

#### Fully connected layers (Figure 3E)

The fully connected neural network layers contained input, hidden and output layers (Figure 2E) with various numbers of neurons (*l*_*r*_). Every neuron in a layer was connected to neurons in the next layer [76]. Fully connected layers received the output of LSTM layers as input. The fully connected layers calculated a weighted sum of LSTM outputs and added a bias term to the outputs. These data were then passed to an activation function (*f*) to define the output for each neuron (Eqs. 8 and 9) [77].

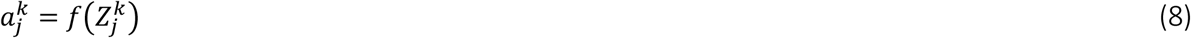

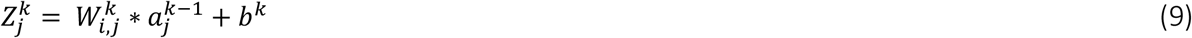

Where *k* ∈ {*1*, …, *n*} and (*i, j*) represent the number of hidden layers and neurons in each pair of subsequent hidden layers (*l*_*r*_, *l*_*r*+1_). The optimized values for these parameters were found via hyperparameter tunning where, *a*^*k*^ is each neuron output where *a*^0^ ∈ {*h*_1_, …, *h*_*m*_} is the LSTM layers output and the input to the fully connected layers and *a*^*n*+1^ is the network output: 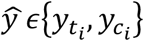 where 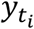 and 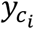 are the outputs for translation and classification tasks, respectively. We first assigned random values to all network parameters *θ*_*t*_; each neuron weight (*W*_*i,j*_) (Figure 3E), bias term (*b*) which is a constant added to calculate the neurons output and other network hyperparameters (the number of hidden layers, the number of neurons for each hidden layer and activation functions for each hidden layer) to start the optimization process for finding the best network infrastructure. Next, we estimated the network errors using mean squared error, MSE (Eq. 10) and cross-entropy loss functions (Eq. 11) to map the translation and classification tasks [54, 78], respectively.

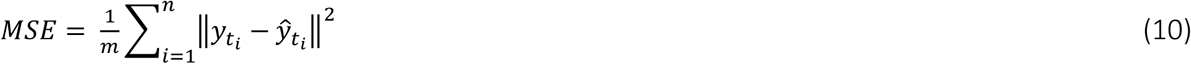

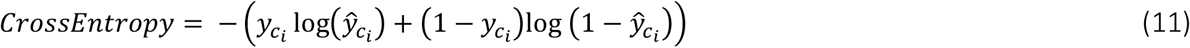

where *m* is the total number of LSTM layers outputs (*h*_*m*_) and 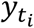 and 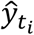 are the simulated and translated adult-CM APs (the network output for translation task). The 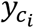 is binary indicator of class labels for iPSC-CM APs (0 for drug-free or 1 for drugged categories) and 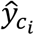 is predicted probability of APs being classified into the discussed classes. We used sum of both loss functions (Eq. 12) to calculate the overall network error (*J*) for both translation and classification tasks during the network training process. We updated network parameters (*θ*_*t*+1_) using adaptive momentum estimation (ADAM) optimization algorithm [79] based on the average gradient of overall loss function with respect to the network parameters for 64 randomly selected simulated AP traces (mini-batch = 64) at each training iteration (Eqs. 13-15).

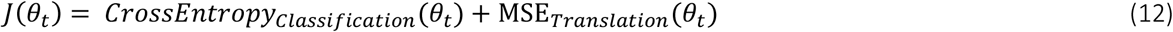

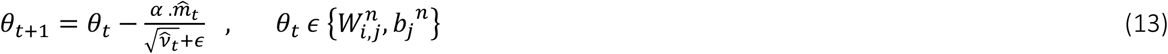

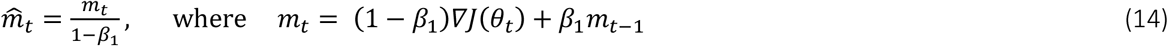

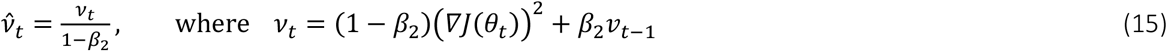

We used a rectified linear unit (ReLu) [80] as activation function in Eq. 8 to calculate the output for each hidden layer neuron at each training iteration. We used dropout regularization [81] to randomly drop neurons with 0.2 probability of elimination along with their connections from the LSTM and fully connected layers during training to reduce the overfitting. We kept updating the network parameters using ADAM optimization algorithm (Eq. 13) to find global minimum of loss function (Eq. 12). We computed the exponential average of the gradient (Eq. 14) as well as the square of the gradient (Eq. 15) for each parameter (θ_*t*_) where *α* is the learning rate equal to 0.001, *β*_1_, *β*_2_ are first and second momentum coefficients equal to *0*.*9* and *0*.*999*, and *ϵ* is a small term equal to *1e*^*-*8^ preventing division by zero.

### Computational workflow (Figure 4)

We first preprocessed iPSC-CM and adult-CM APs by applying a digital forward and backward data filtering technique [56] and normalizing the AP values for more efficient training process. Next, we split the preprocessed data in 70:10:20 ratio into training, validation and test data sets, respectively, and implemented the network architecture using Pytorch [82]. During the training process the multitask network received iPSC-CM AP time course data as inputs and predicted adult-CM AP time courses. The network also received the category (drug-free and drugged) of the iPSC-CM AP data. The network next calculated the MSE (Eq. 10) between predicted AP waveforms and the expected waveforms for adult-CM APs. It also calculated cross-entropy (Eq. 11) between the predicted category for the iPSC-CM AP and the expected value. The cross-entropy was added to the calculated MSE to determine the total loss for training. The ADAM optimization algorithm was then used to update the network weights and bias terms.

We performed updating the network parameters (Eq. 13) and monitored the network performance for the training and validation data sets until the point at which the network performance on the training data set began to degrade compared to the validation dataset. This process was used to identify the optimal number of iterations (epochs = 300) for the training process. The last trained network was designated as the best possible model to perform both translation and classification tasks. We then used a holdout test dataset and calculated MSE (Eq. 10), R^2^_score (Eqs. 16-17 below) and the error in prediction for adult-CM *APD*_*90*_ as evaluation metrics to assess the performance of the network for translation task and the area under the receiver operating characteristic (AUROC) curve, accuracy, recall and precision to measure capability of network for classification task as described below. The network codes have been made publicly available at Clancy lab Github. (https://github.com/ClancyLabUCD/Multitask_network)

### Evaluation metrics for the translation and classification tasks

As we discussed, we used MSE and cross-entropy loss functions for performance evaluation of translation and classification tasks. In addition to MSE, we computed R^2^_score [57] (Eqs. 16,17) to measure how close the translated adult-CM AP 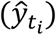 was to the expected simulated adult-CM AP 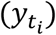. We compared the histogram distribution of simulated and translated adult-CM APD_90_ values and the error in *APD*_*90*_ prediction to assess the accuracy of network prediction.

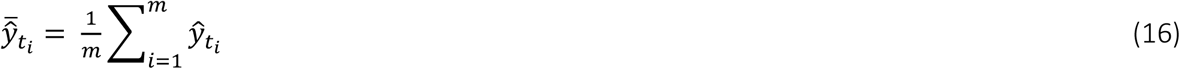

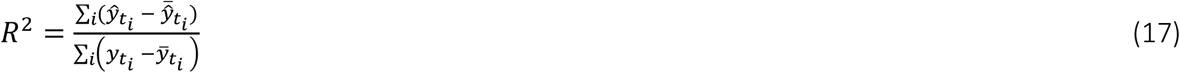

We used AUROC to measure the capability of the model to distinguish between drug-free and drugged iPSC-CM APs [58]. AUROC is the area under the Receiver Operating Characteristic (ROC) curve, which is a plot of the false positive rate (FPR), the probability that the network classified drug-free iPSC-CM APs into drugged categories (FP) (Eq. 18) versus the true positive rate (TPR) or recall, the probability that the network correctly classified drugged iPSC-CM APs into drugged category (TP) (Eq. 19). AUROC close to 1 indicated a model with a desirable measure of separability, while a poor model had AUROC near 0, which means that it had poor separability.

In addition, we used recall, accuracy, and precision to describe the performance of the network for the classification task [13], where the accuracy and precision indicated the proportion of all correct, TP + true negatives (TN), i.e., predicted drug-free APs (Eq. 20) and correct positive identifications (Eq. 21). False negatives (FN) in Eqs. 19-20 were the total number of drugged iPSC-CM APs classified as drug-free.

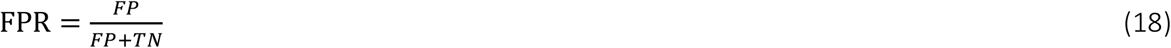

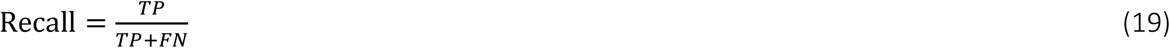

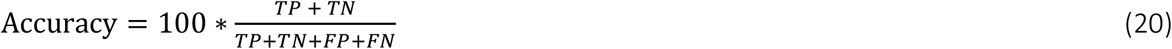

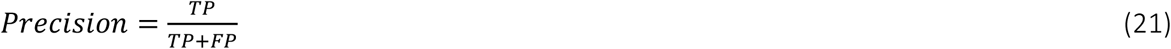

## Notes

https://github.com/ClancyLabUCD/Multitask_network

### Competing Interest Statement

The authors have declared no competing interest.

## References

1. Shaheen, N., et al., Human induced pluripotent stem cell -derived cardiac cell sheets expressing genetically encoded voltage indicator for pharmacological and arrhythmia studies. Stem cell reports, 2018. 10(6): p. 1879–1894. https://doi.org/10.1016/j.stemcr.2018.04.006

2. Leyton-Mange, J.S., et al., Rapid cellular phenotyping of human pluripotent stem cell - derived cardiomyocytes using a genetically encoded fluorescent voltage sensor. Stem cell reports, 2014. 2(2): p. 163–170. https://doi.org/10.1016/j.stemcr.2014.01.003

3. Sun, N., et al., Patient-specific induced pluripotent stem cells as a model for familial dilated cardiomyopathy. Sci Transl Med, 2012. 4(130): p. 130ra47. https://doi.org/10.1126/scitranslmed.3003552

4. Lan, F., et al., Abnormal calcium handling properties underlie familial hypertrophic cardiomyopathy pathology in patient -specific induced pluripotent stem cells. Cell Stem Cell, 2013. 12(1): p. 101–13. https://doi.org/10.1016/j.stem.2012.10.010

5. Burridge, P.W., et al., Human induced pluripotent stem cell -derived cardiomyocytes recapitulate the predilection of breast cancer patients to doxorubicin-induced cardiotoxicity. Nat Med, 2016. 22(5): p. 547–56. https://doi.org/10.1038/nm.4087

6. Doss, M.X. and A. Sachinidis, Current challenges of iPSC-based disease modeling and therapeutic implications. Cells, 2019. 8(5): p. 403. https://doi.org/10.3390/cells8050403

7. Collins, T.A., M.G. Rolf, and A. Pointon, Current and future approaches to nonclinical cardiovascular safety assessment. Drug Discovery Today, 2020. https://doi.org/10.1016/j.drudis.2020.03.011

8. Wu, J.C., et al., Towards precision medicine with human iPSCs for cardiac channelopathies. Circulation research, 2019. 125(6): p. 653–658. https://doi.org/10.1161/CIRCRESAHA.119.315209

9. Sayed, N., C. Liu, and J.C. Wu, Translation of human-induced pluripotent stem cells: from clinical trial in a dis h to precision medicine. Journal of the American College of Cardiology, 2016. 67(18): p. 2161-2176. http://dx.doi.org/10.1016/j.jacc.2016.01.083

10. Matsa, E., J.H. Ahrens, and J.C. Wu, Human induced pluripotent stem cells as a platform for personalized and precision cardiovascular medicine. Physiological Reviews, 2016. 96(3): p. 1093–1126. https://doi.org/10.1152/physrev.00036.2015

11. Tveito, A., et al., Inversion and computational maturation of drug response using human stem cell derived cardiomyocytes in microphysiological systems. Scientific reports, 2018. 8(1): p. 1–14. https://doi.org/10.1038/s41598-018-35858-7

12. Tveito, A., et al., Computational translation of drug effects from animal experiments to human ventricular myocytes. Scientific Reports, 2020. 10(1): p. 1–11. https://doi.org/10.1038/s41598-020-66910-0

13. Sube, R. and E.A. Ertel, Cardiomyocytes Derived from Human Induced Pluripotent Stem Cells: An In-Vitro Model to Predict Cardiac Effects of Drugs. Journal of Biomedical Science and Engineering, 2017. 10(11): p. 527. https://doi.org/10.4236/jbise.2017.1011040

14. Navarrete, E.G., et al., Screening drug-induced arrhythmia using human induced pluripotent stem cell –derived cardiomyocytes and low-impedance microelectrode arrays. Circulation, 2013. 128(11_suppl_1): p. S3–S13. https://doi.org/10.1161/CIRCULATIONAHA.112.000570

15. Lieu, D.K., et al., Mechanism-based facilitated maturation of human pluripotent stem cell-derived cardiomyocytes. Circ Arrhythm Electrophysiol, 2013. 6(1): p. 191–201. https://doi.org/10.1161/CIRCEP.111.973420

16. Veerman, C.C., et al., Immaturity of human stem-cell-derived cardiomyocytes in culture: fatal flaw or soluble problem? Stem Cells Dev, 2015. 24(9): p. 1035–52. https://doi.org/10.1089/scd.2014.0533

17. Tu, C., B.S. Chao, and J.C. Wu, Strategies for Improving the Maturity of Human Induced Pluripotent Stem Cell -Derived Cardiomyocytes. Circ Res, 2018. 123(5): p. 512–514. https://doi.org/10.1161/CIRCRESAHA.118.313472

18. Blinova, K., et al., International multisite study of human -induced pluripotent stem cell - derived cardiomyocytes for drug proarrhythmic potential assessment. Cell reports, 2018. 24(13): p. 3582–3592. https://doi.org/10.1016/j.celrep.2018.08.079

19. Sala, L., M. Bellin, and C.L. Mummery, Integrating cardiomyocytes from human pluripotent stem cells in safety pharmacology: has the time come? British journal of pharmacology, 2017. 174(21): p. 3749–3765. https://doi.org/10.1111/bph.13577

20. Gong, J.Q. and E.A. Sobie, Population-based mechanistic modeling allows for quantitative predictions of drug responses across cell types. NPJ systems biology and applications, 2018. 4(1): p. 1–11. https://doi.org/10.1038/s41540-018-0047-2

21. de Korte, T., et al., Unlocking personalized biomedicine and drug discovery with human induced pluripotent stem cell –derived Cardiomyocytes: fit for purpose or forever elusive? Annual Review of Pharmacology and Toxicology, 2020. 60: p. 529–551. https://doi.org/10.1146/annurev-pharmtox-010919-023309

22. Koivumäki, J.T., et al., Structural immaturity of human iPSC -derived cardiomyocytes: in silico investigation of effects on function and disease modeling. Frontiers in physiology, 2018. 9: p. 80. https://doi.org/10.3389/fphys.2018.00080

23. Alhusseini, M.I., et al., Machine Learning to Classify Intracardiac Electrical Patterns During Atrial Fibrillation: Machine Learning of Atrial Fibrillation. Circulation: Arrhythmia and Electrophysiology, 2020. 13(8): p. e008160. https://doi.org/10.1161/CIRCEP.119.008160

24. Rogers, A.J., et al., Machine Learned Cellular Phenotypes Predict Outcome in Ischemic Cardiomyopathy. Circulation Research, 2020. https://doi.org/10.1161/CIRCRESAHA.120.317345

25. Sevakula, R.K., et al., State-of-the-Art machine learning techniques aiming to improve patient outcomes pertaining to the cardiovascular system. Journal of the American Heart Association, 2020. 9(4): p. e013924. https://doi.org/10.1161/JAHA.119.013924

26. Jin, Z., et al. HeartToGo: a personalized medicine technology for cardiovascular disease prevention and detection. in 2009 IEEE/NIH Life Science Systems and Applications Workshop. 2009. IEEE. https://doi.org/10.1109/LISSA.2009.4906714

27. Trayanova, N.A., D.M. Popescu, and J.K. Shade, Machine Learning in Arrhythmia and Electrophysiology. Circulation Research, 2021. 128(4): p. 544–566. https://doi.org/10.1161/CIRCRESAHA.120.317872

28. Hochreiter, S. and J. Schmidhuber, Long short-term memory. Neural computation, 1997. 9(8): p. 1735–1780. https://doi.org/10.1162/neco.1997.9.8.1735

29. Guo, A., et al., Predicting cardiovascular health trajectories in time -series electronic health records with LSTM models. BMC Medical Informatics and Decision Making, 2021. 21(1): p. 1–10. https://doi.org/10.1186/s12911-020-01345-1

30. Shi, K., et al., Contactless analysis of heart rate variability during cold pressor test using radar interferometry and bidirectional LSTM networks. Scientific reports, 2021. 11(1): p. 1–13. https://doi.org/10.1038/s41598-021-81101-1

31. Picon, A., et al., Mixed convolutional and long short -term memory network for the detection of lethal ventricular arrhythmia. PloS one, 2019. 14(5): p. e0216756. https://doi.org/10.1371/journal.pone.0216756

32. Ballinger, B., et al. DeepHeart: semi-supervised sequence learning for cardiovascular risk prediction. in Thirty-Second AAAI Conference on Artificial Intelligence. 2018.

33. He, R., et al., Automatic cardiac arrhythmia classification using combination of deep residual network and bidirectional LSTM. IEEE Access, 2019. 7: p. 102119–102135. https://doi.org/10.1109/ACCESS.2019.2931500

34. Hou, B., et al., LSTM Based Auto-Encoder Model for ECG Arrhythmias Classification. IEEE Transactions on Instrumentation and Measurement, 2019. https://doi.org/10.1109/TIM.2019.2910342

35. Warrick, p. and M.N. Homsi. Cardiac arrhythmia detection from ECG combining convolutional and long short -term memory networks. in 2017 Computing in Cardiology (CinC). 2017. IEEE. https://doi.org/10.22489/CinC.2017.161-460

36. Oh, S.L., et al., Automated diagnosis of arrhythmia using combination of CNN and LSTM techniques with variable length heart beats. Computers in biology and medicine, 2018. 102: p. 278–287. https://doi.org/10.1016/j.compbiomed.2018.06.002

37. Chen, C., et al., Automated arrhythmia classification based on a combination network of CNN and LSTM. Biomedical Signal Processing and Control, 2020. 57: p. 101819. https://doi.org/10.1016/j.bspc.2019.101819

38. Wang, L. and X. Zhou, Detection of congestive heart failure based on LSTM -based deep network via short-term RR intervals. Sensors, 2019. 19(7): p. 1502. https://doi.org/10.3390/s19071502

39. Bian, M., et al. An accurate lstm based video heart rate estimation method. in Chinese Conference on Pattern Recognition and Computer Vision (PRCV) .2019.Springer. https://doi.org/10.1007/978-3-030-31726-3_35

40. Maragatham, G. and S. Devi, LSTM model for prediction of heart failure in big data. Journal of medical systems, 2019. 43(5): p. 1–13. https://doi.org/10.1007/s10916-019-1243-3

41. Yildirim, O., et al., A new approach for arrhythmia classification using deep coded features and LSTM networks. Computer methods and programs in biomedicine, 2019. 176: p. 121–133. https://doi.org/10.1016/j.cmpb.2019.05.004

42. Wang, E.K., X. Zhang, and L. Pan, Automatic classification of CAD ECG signals with SDAE and bidirectional long short -term network. IEEE Access, 2019. 7: p. 182873–182880. https://doi.org/10.1109/ACCESS.2019.2936525

43. Martis, R.J., et al., Application of higher order cumulant features for cardiac health diagnosis using ECG signals. International journal of neural systems, 2013. 23(04): p. 1350014. https://doi.org/10.1142/S0129065713500147

44. Liu, F., et al. A LSTM and CNN based assemble neural network framework for arrhythmias classification. in ICASSP 2019-2019 IEEE International Conference on Acoustics, Speech and Signal Processing (ICASSP). 2019. IEEE. https://doi.org/10.1109/ICASSP.2019.8682299

45. Yildirim, Ö., A novel wavelet sequence based on deep bidirectional LSTM network model for ECG signal classification. Computers in biology and medicine, 2018. 96: p. 189–202. https://doi.org/10.1016/j.compbiomed.2018.03.016

46. Yang, P.-C., et al., A computational pipeline to predict cardiotoxicity: From the atom to the rhythm. Circulation research, 2020. 126(8): p. 947–964. https://doi.org/10.1161/CIRCRESAHA.119.316404

47. Cai, C., et al., Deep learning-based prediction of drug-induced cardiotoxicity. Journal of chemical information and modeling, 2019. 59(3): p. 1073–1084. https://doi.org/10.1021/acs.jcim.8b00769

48. Zhang, Y., et al., Prediction of hERG K+ channel blockage using deep neural networks. Chemical Biology & Drug Design, 2019. 94(5): p. 1973–1985. https://doi.org/10.1111/cbdd.13600

49. Dickson, C.J., C. Velez-Vega, and J.S. Duca, Revealing molecular determinants of hERG blocker and activator binding. Journal of chemical information and modeling, 2019. 60(1): p. 192–203. https://doi.org/10.1021/acs.jcim.9b00773

50. Ryu, J.Y., et al., DeepHIT: a deep learning framework for prediction of hERG -induced cardiotoxicity. Bioinformatics, 2020. 36(10): p. 3049–3055. https://doi.org/10.1093/bioinformatics/btaa075

51. Li, Z., et al., General principles for the validation of proarrhythmia risk prediction models: an extension of the CiPA in silico strategy. Clinical Pharmacology & Therapeutics, 2020. 107(1): p. 102–111. https://doi.org/10.1002/cpt.1647

52. Kernik, D.C., et al., A computational model of induced pluripotent stem-cell derived cardiomyocytes incorporating experimental variability from multiple data sources. The Journal of physiology, 2019. https://doi.org/10.1113/JP277724

53. O’Hara, T., et al., Simulation of the undiseased human cardiac ventricular action potential: model formulation and experimental validation. PLoS computational biology, 2011. 7(5): p. e1002061. https://doi.org/10.1371/journal.pcbi.1002061

54. Goodfellow, I., Y. Bengio, and A. Courville, Deep learning. 2016: MIT press. https://doi.org/10.4258/hir.2016.22.4.351

55. Tanskanen, A.J. and L.H. Alvarez, Voltage noise influences action potential duration in cardiac myocytes. Mathematical biosciences, 2007. 208(1): p. 125–146. https://doi.org/10.1016/j.mbs.2006.09.023

56. Gustafsson, F., Determining the initial states in forward -backward filtering. IEEE Transactions on signal processing, 1996. 44(4): p. 988–992. https://doi.org/10.1109/78.492552

57. Devore, J.L., Probability and Statistics for Engineering and the Sciences. 2011: Cengage learning.

58. Fawcett, T., An introduction to ROC analysis. Pattern recognition letters, 2006. 27(8): p. 861–874. https://doi.org/10.1016/j.patrec.2005.10.010

59. Powers, D.M., Evaluation: from precision, recall and F-measure to ROC, informedness, markedness and correlation. 2011.

60. LeCun, Y., J. Denker, and S. Solla, Optimal brain damage. Advances in neural information processing systems, 1989 2: p. 598–605.

61. Reale, R.A., J.F. Brugge, and J.C. Chan, Maps of auditory cortex in cats reared after unilateral cochlear ablation in the neonatal period. Developmental Brain Research, 1987. 34(2): p. 281–290. https://doi.org/10.1016/0165-3806(87)90215-X

62. Yang, P.C., et al., A computational modelling approach combined with cellular electrophysiology data provides insights into the therapeutic benefit of targeting the late Na+ current. The Journal of physiology, 2015. 593(6): p. 1429–1442. https://doi.org/10.1113/jphysiol.2014.279554

63. Fabbri, A., et al., Required GK1 to suppress automaticity of iPSC -CMs depends strongly on IK1 model structure. Biophysical Journal, 2019. 117(12): p. 2303–2315. https://doi.org/10.1016/j.bpj.2019.08.040

64. Casini, S., A.O. Verkerk, and C.A. Remme, Human iPSC-derived cardiomyocytes for investigation of disease mechanisms and therapeutic strategies in inherited arrhythmia syndromes: strengths and limitations. Cardiovascular drugs and therapy, 2017. 31(3): p. 325–344. https://doi.org/10.1007/s10557-017-6735-0

65. Goversen, B., et al., The immature electrophysiological phenotype of iPSC -CMs still hampers in vitro drug screening: Special focus on IK1. Pharmacology & therapeutics, 2018 183: p. 127–136. https://doi.org/10.1016/j.pharmthera.2017.10.001

66. Knollmann, B.C., Induced pluripotent stem cell –derived cardiomyocytes: Boutique science or valuable arrhythmia model? Circulation research, 2013. 112(6): p. 969–976. https://doi.org/10.1161/CIRCRESAHA.112.300567

67. Sinnecker, D., et al., Induced pluripotent stem cell -derived cardiomyocytes: a versatile tool for arrhythmia research. Circulation Research, 2013. 112(6): p. 961–968. https://doi.org/10.1161/CIRCRESAHA.112.268623

68. Blinova, K., et al., Comprehensive translational assessment of human -induced pluripotent stem cell derived cardiomyocytes for evaluating drug -induced arrhythmias. Toxicological Sciences, 2017. 155(1): p. 234–247. https://doi.org/10.1093/toxsci/kfw200

69. Kernik, D.C., et al., A computational model of induced pluripotent stem -cell derived cardiomyocytes for high throughput risk stratification of KCNQ1 genetic variants. PLOS Computational Biology, 2020. 16(8): p. e1008109. https://doi.org/10.1371/journal.pcbi.1008109

70. Yoshinaga, D., et al., Phenotype-based high-throughput classification of long QT syndrome subtypes using human induced pluripotent stem cells. Stem cell reports, 2019. 13(2): p. 394–404. https://doi.org/10.1016/j.stemcr.2019.06.007

71. Atkinson, K.E., An introduction to numerical analysis. 2008: John wiley & sons.

72. Yang, P.C., et al., A Computational Pipeline to Predict Cardiotoxicity: From the Atom to the Rhythm. Circ Res, 2020. 126(8): p. 947–964. https://doi.org/10.1161/CIRCRESAHA.119.316404

73. Li, M., et al., Overexpression of KCNJ2 in induced pluripotent stem cell -derived cardiomyocytes for the assessment of QT-prolonging drugs. Journal of Pharmacological Sciences, 2017. 134(2): p. 75–85. https://doi.org/10.1016/j.jphs.2017.05.004

74. Cheng, J., L. Dong, and M. Lapata, Long short-term memory-networks for machine reading. arXiv preprint arXiv:1601.06733, 2 016.

75. Olah, C., Understanding LSTM Networks. Aug. 2015. URL https://colah.github.io/posts/2015-08-Understanding-LSTMs, 2017.

76. Krogh, A., What are artificial neural networks? Nature biotechnology, 2008. 26(2): p. 195–197. https://doi.org/10.1038/nbt1386

77. Carugo, O., F. Eisenhaber, and Carugo, Data mining techniques for the life sciences . Vol. 609. 2010: Springer. https://doi.org/10.1007/978-1-4939-3572-7

78. Murphy, K.P., Machine learning: a probabilistic perspective. 2012: MIT press.

79. Kingma, D.p. and J. Ba, Adam: A method for stochastic optimization. arXiv preprint arXiv:1412.6980, 2014.

80. Glorot, X., A. Bordes, and Y. Bengio. Deep sparse rectifier neural networks. in Proceedings of the fourteenth international conference on artificial intelligence and statistics. 2011. JMLR Workshop and Conference Proceedings.

81. Zaremba, W., I. Sutskever, and O. Vinyals, Recurrent neural network regularization. arXiv preprint arXiv:1409.2329, 2014.

82. Ketkar, N., Introduction to pytorch, in Deep learning with python. 2017, Springer. p. 195–208. https://doi.org/10.1007/978-1-4842-2766-4_12

